# Nicotinic and Cholinergic Modulation of Reward Prediction Error Computations in the Ventral Tegmental Area: a Minimal Circuit Model

**DOI:** 10.1101/423806

**Authors:** Nicolas Deperrois, Boris Gutkin

## Abstract

Dopamine (DA) neurons in the ventral tegmental area (VTA) are thought to encode reward prediction errors (RPE) by comparing actual and expected rewards. In recent years, much work has been done to identify how the brain uses and computes this signal.While several lines of evidence suggest the the interplay of he DA and the inhibitory interneurons in the VTA implements the RPE computaiton, it still remains unclear how the DA neurons learn key quantities, for example the amplitude and the timing of primary rewards during conditioning tasks. Furthermore, exogenous nicotine and endogenous acetylcholine, acting on both VTA DA and GABA (*γ* - aminobutyric acid) neurons via nicotinic-acetylcholine receptors (nAChRs), also likely affect these computations. To explore the potential circuit-level mechanisms for RPE computations during classical-conditioning tasks, we developed a minimal computational model of the VTA circuitry. The model was designed to account for several reward-related properties of VTA afferents and recent findings on VTA GABA neuron dynamics during conditioning.

With our minimal model, we showed that the RPE can be learned by a two-speed process computing reward timing and magnitude. Including a model of nAChR-mediated currents in the VTA DA-GABA circuit, we also showed that nicotine should reduce the acetylcholine action on the VTA GABA neurons by receptor desensitization and therefore potentially boost the DA responses to reward information. Together, our results delineate the mechanisms by which RPE are computed in the brain, and suggest a hypothesis on nicotine-mediated effects on reward-related perception and decision-making.

## 1 Introduction

To adapt to their environment, animals constantly compare their predictions with new environmental outcomes (rewards, punishments, etc). The difference between prediction and outcome is the prediction error, which in turn can serve as a teaching signal to allow the animal to update its predictions and render previously neutral stimuli predictive of rewards into reinforcers of behavior. Particularly, the dopamine (DA) neuron activity in the Ventral Tegmental Area (VTA) have been shown to encode the reward prediction error (RPE), or the difference between the actual reward the animal receives and the expected reward (Schultz et al., 1997; Schultz, 1998; Bayer and Glimcher, 2005; Day and Carelli, 2007; Enomoto et al., 2011; Matsumoto and Hikosaka, 2009; Eshel et al., 2015; Keiflin and Janak, 2015). During, for example, classical conditioning with appetitive rewards, unexpected rewards elicit strong transient increases in VTA DA neuron activity, but as a cue fully predicts the reward, the same reward produces little or no DA neurons response. Finally, after learning, if the reward is omitted, DA neurons pause their firing at the moment reward is expected (Schultz et al., 1997; Schultz, 1998; Keiflin and Janak, 2015; Watabe-Uchida et al., 2017). Thus DA neurons should either receive or compute the RPE. While several lines of evidence have pointed towards the RPE being computed by the VTA local circuitry, exactly how this is done vis-a-vis the inputs and how this computation is modulated by the endogenous acetylcholine and the endogenous substances that affect the VTA, *e.g*. nicotine, remains to be defined. Here we proceed to address these questions using a minimal computational modelling methodology.

In order to compute the RPE, the VTA should receive the relevant information from it inputs. Intuitively, distinct biological inputs to the VTA must differentially encode actual and expected rewards that are finally subtracted by a downstream target, the VTA DA neurons. For the last two decades, a great amount of experimental studies depicted which brain areas send this information to the VTA. Notably, a subpopulation of pedunculopontine tegmental nucleus (PPTg) has been found to send the actual reward signal to dopamine neurons (Kobayashi and Okada, 2007; Okada et al., 2009; Keiflin and Janak, 2015), while other studies showed that the prefrontal cortex (PFC) and the nucleus accumbens (NAc) respond to the predictive cue (Keiflin and Janak, 2015; Oyama et al., 2015; Funahashi, 2006; Connor and Gould, 2016; Le Merre et al., 2018), highly depending on VTA DA feedback projections in the PFC (Puig et al., 2014; Popescu et al., 2016) and the NAc (Yagishita et al., 2014; Keiflin and Janak, 2015; Fisher et al., 2017). However, how each of these signals are integrated by VTA DA neurons during classical-conditioning remains elusive.

Recently, VTA GABA neurons were shown to encode reward expectation with a persistent cue response proportional to the expected reward (Cohen et al., 2012; Eshel et al., 2015; Tian et al., 2016). Additionally, selectively exciting and inhibiting VTA GABA neurons during a classical-conditioning task, Eshel et al. (2015) revealed that these neurons are likely source of the substraction operation, contributing to the inhibitory expectation signal in the RPE computation by DA neurons.

Furthermore, the presence of nicotinic acetylcholine receptors (nAChRs) in the VTA (Pontieri et al., 1996; Maskos et al., 2005; Changeux, 2010; Faure et al., 2014) provides a potential common route for acetylcholine (ACh) and nicotine (Nic) in modulating dopamine activity during a Pavlovian-conditioning task. Particularly, the high-affinity α4β2 subunit-containing nAChRs desensitizing relatively slowly (≃ sec) and located post-synaptically on VTA DA and GABA neurons have been shown to have the most prominent role in nicotine-induced DAergic bursting activity and self-administration, as suggested by mouse knock-out experiments (Maskos et al., 2005; Changeux, 2010; Faure et al., 2014) and recent direct optogenetic modulation of these somatic receptors (Durand-de Cuttoli et al., 2018).

We have previously developed and validated a population level circuit dynamics model (Graupner et al., 2013; Tolu et al., 2013; Maex et al., 2014; Dumont et al., 2018) of the influence nicotine and Ach interplay may have on the VTA dopamine cell activity. Using this model we showed that Nic action on α4β2 could result in either direct stimulation or disinhibition of DA neurons. The latter scenario suggests that relatively low nicotine concentrations (^~^500 nM) during and after smoking preferentially desensitize α4β2 nAChRs on GABA neurons (Fiorillo et al., 2008). The endogenous cholinergic drive to GABA neurons would then decrease, resulting in decreased GABA neurons activity, and finally a disinhibition of DA neurons as confirmed *in vitro* (Mansvelder et al., 2002) and suggested by Graupner et al. (2013); Tolu et al. (2013); Maex et al. (2014); Dumont et al. (2018) modeling work. Interestingly, this scenario requires that the high affinity nAChRs are in a pre-activated state, so that nicotine can desensitize them, which in turn implies a sufficiently high ambient cholinergic tone in the VTA. However, when the ACh tone is not sufficient, in this GABA-nAChR scenario, nicotine would lead to a significant inhibition of the DA neurons. Furthermore, a recent study showed that optogenetic inhibition of PPTg cholinergic fibers inhibit only the VTA non-DA neurons (Yau et al., 2016), suggesting that ACh acts preferentially on VTA GABA neurons. However, the effects of Nic and ACh on dopamine responses to rewards via α4β2-nAChRs desensitization during classical-conditioning have remained elusive.

In addition to the above issues, a non-trivial question comes from the timing structure of the conditioning tasks. Typically, the reward to be consumed is delivered after a temporal delay after the conditioning cue, which begs important related questions: how is the reward information transferred from the reward-delivery time to the earlier reward-predictive stimulus and how does the brain compute the precise timing of reward? In other words, how is the relative co-timing of the reward and the reinforcer learned in the brain? These issues generate further lines of enquiry on how this learning process may be altered by nicotine. In order to start clarifying the possible neural mechanisms underlying the observed RPE-like activity in DA neurons, we propose here a simple neuro-computational model inspired from Graupner et al. (2013), incorporating the mean dynamics of four neuron populations: the prefrontal cortex (PFC), the pedunculopontine tegmental nucleus (PPTg), the VTA dopamine and GABA neurons. Taking into account recent neurobiological data, particularly showing the activity of VTA GABA neurons (Cohen et al., 2012; Eshel et al., 2015) during classical-conditioning, we qualitatively and quantitively reproduce several aspects of a Pavlovian-conditioning task - which we take as a paradigmatic example of reward-based conditioning - such as the phasic components of dopaminergic activation with respect to reward magnitude, omission and timing, the working-memory activity in the PFC, the response of the PPTg to primary rewards, and the dopamine-induced plasticity in cortical and corticostriatal synapses. Finally, we qualitatively assessed the potential effects of Nic-induced desensitization of GABA α4β2-nAChRs, leading to a disinhibition of DA burst-response to rewarding events.

## 2 Methods: Computational Model and Simulated Behavioral Tasks

In order to examine the effects of nicotine on VTA activity during classical-conditioning, we built a neural population model of the VTA and its afferent inputs inspired the mean-field approach from Graupner et al. (2013). This model incorporates the DA and GABA neuronal populations in the VTA and their glutamatergic and cholinergic afferents from the PFC and the PPTg (Fig. 1). Based on recent neurobiological data, we propose a model for the activity of the PFC and PPTg inputs during classical-conditioning contributing to the observed VTA GABA and DA activity. Additionally, the activation and desensitization dynamics of the nAChR-mediated currents in response to Nic and ACh were described by a 4-state model taken from Graupner et al. (2013).

**Figure 1.**
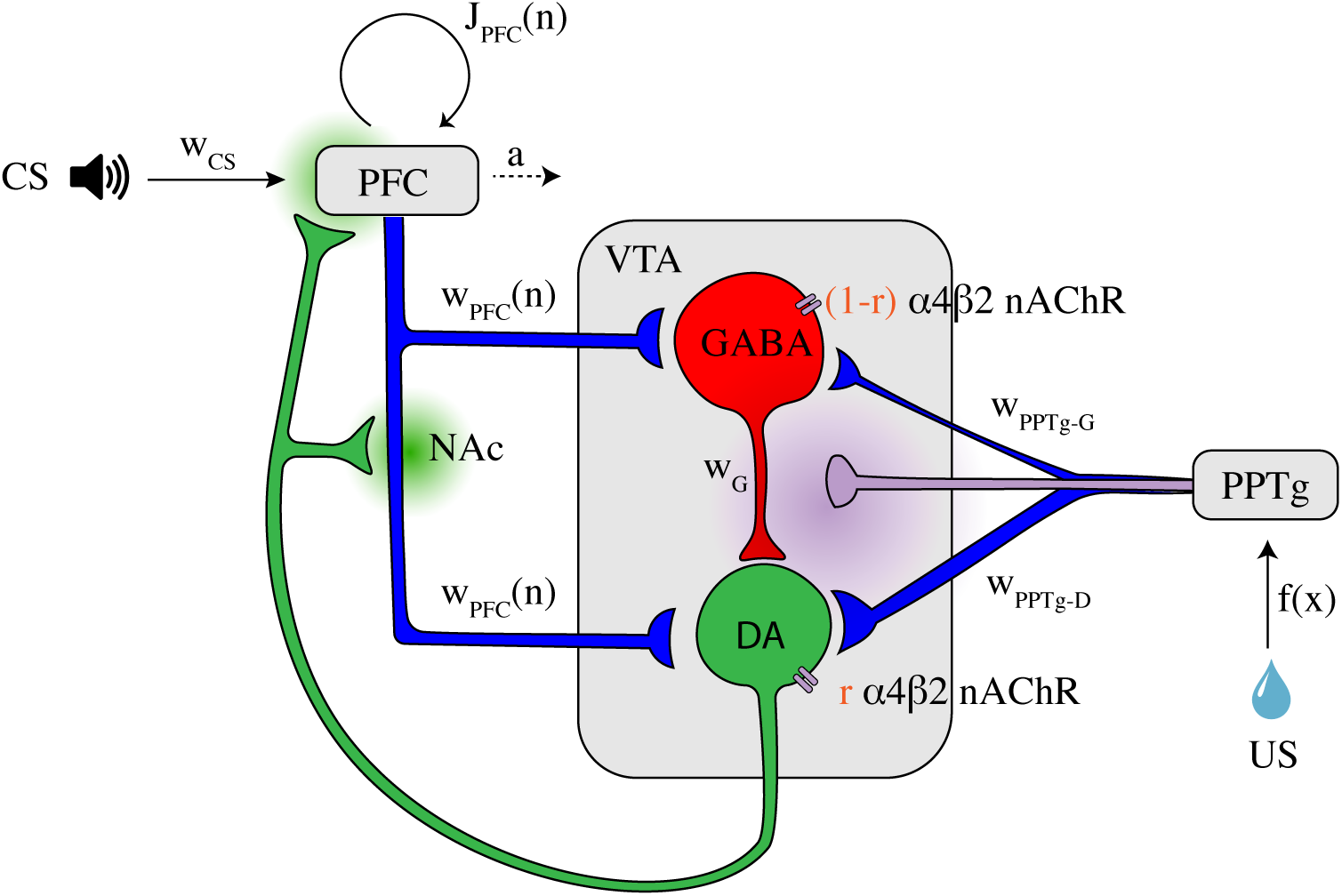
Illustration of the VTA circuit and neural dynamics of each area during learning of a pavlovian-conditioning task Afferents inputs and circuitry of the ventral tegmental area (VTA). The GABA neuron population (red) inhibits locally the DA neuron population (green). This local circuit receives excitatory glutamatergic input (blue axons) from the corticostriatal pathway and the pedunculopontine tegmental nucleus (PPTg). The PPTg furthermore furnishes cholinergic projections (purple axon) to the VTA neurons (α4β2 nAChRs). *r* is the parameter to change continuously the dominant site of α4β2 nAChR action. Dopaminergic efferents (green axon) project, amongst others, to the nucleus accumbens (NAc) and the prefrontal cortex (PFC) and modulates cortico-striatal projections *w*_PFC_ and PFC recurrent excitation *J*_PFC_ weights. The PFC integrates CS (tone) information, while the PPTg respond phasically to the water reward itself (US). Dopamine and acetylcholine outflows are represented by green and purple shaded areas, respectively. All parameters and description are summarized in Table 1.

**Table 1.** Model parameters

| Parameter | Description | Value | Reference |
| --- | --- | --- | --- |
| **$\alpha 4\beta 2$-nAChR** |  |  | (Graupner et al., 2013) |
| $EC_{50}$ | half-maximum conc. of activation (ACh) | 30 μM | |
| $\alpha$ | potency of Nic to evoke response | 3 | |
| $n_a$ | Hill coefficient of activation | 1.05 | |
| $IC_{50}$ | half-maximum conc. of desensitization by Nic | 0.061 μM | |
| $n_d$ | Hill coefficient of desensitization | 0.5 | |
| $\tau_a$ | activation time constant | 5 msec | |
| $K_\tau$ | half-maximum conc. of desensitization time const. | 0.11 μM | |
| $n_\tau$ | Hill coefficient of desensitization time constant | 3 | |
| $\tau_{\max}$ | maximal desensitization time constant | 10 min | |
| $\tau_0$ | minimal desensitization time constant | 500 msec | |
| **Network** |  |  |  |
| $x$ | reward size | 1-20 μL | (Eshel et al., 2015) |
| $w_{CS}$ | strength of CS signal | 8 | here |
| $\tau_D$ | membrane time constant of DA population | 30 ms | (Graupner et al., 2013) |
| $\tau_G$ | membrane time constant of GABA population | 30 ms | (Graupner et al., 2013) |
| $\tau_{PFC}$ | membrane time constant of PFC population | 100 ms | (Gerstner et al., 2014) |
| $\tau_a$ | adaptation time constant | 1000 ms | (Gerstner et al., 2014) |
| $\tau_{PPTg}$ | membrane time constant of PPTg population | 80 ms | (Okada et al., 2009) |
| $w_G$ | strength of GABA input to DA | 1 | (Graupner et al., 2013) |
| $w_{PFC}$ | strength of PFC input to GABA and DA | variable | |
| $J_{PFC}$ | strength of PFC recurrent connections | variable | |
| $w_{PPT-D}$ | strength of PPTg Glu input to DA | 0.8 | (Yoo et al., 2017) |
| $w_{PPT-G}$ | strength of PPTg Glu input to GABA | 0.2 | (Yoo et al., 2017) |
| $w_{\alpha 4\beta 2}$ | strength of nAChR activation | 15 | here |
| $w_{ACh}$ | maximal ACh conc. from PPTg | 1 μM | (Graupner et al., 2013) |
| $B_D$ | baseline firing rate of DA (without input) | 18 Hz | (Eshel et al., 2015) |
| $B_G$ | baseline firing rate of GABA | 14 Hz | (Eshel et al., 2015) |
| $B_{PPTg}$ | baseline firing rate of PPTg | 2 Hz | (Okada et al., 2009) |
| $r$ | balance of $\alpha 4\beta 2$ nAChRs | 0.2 | (Mansvelder et al., 2002) |
| $c$ | strength of adaptation in PFC population | 0.6 | here |
| $\alpha_P$ | learning rate of PFC recurrent weight | 0.2 | here |
| $\alpha_S$ | learning rate of cortico-striatal weight | 0.0005 | here |

### 2.1 Mean-field description of VTA neurons and their afferents

First, the model from Graupner et al. (2013) describing the dynamics of VTA neuron populations and the effects of Nic and ACh on nAChRs was re-implemented with several quantitative modifications according to experimental data.

The temporal dynamics of the average activities of DA and GABA neurons in the VTA taken from Graupner et al. (2013) are described by the following equations:

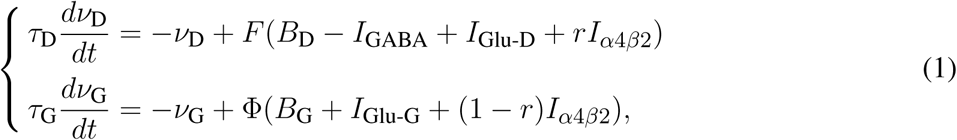

where *ν*_D_ and *ν*_G_ are the mean firing rates of the DA and GABAergic neuron populations, respectively. *τ*_D_ = 30 ms and *τ*_G_ = 30 ms are the membrane time constants of both neuron populations specifying how quickly the neurons integrate input changes. *I*_Glu_ characterize the excitatory inputs from PFC and PPTg mediated by glutamate receptors. *I*_*α*4*β*2_ represent the excitatory input mediated by α4β2-containing nAChRs, activated by PPTg ACh input and Nic. *I*_GABA_ is the local feed-forward inhibitory input to DA neurons emanating from VTA GABA neurons. *B*_D_ = 18 and *B*_G_ = 14 are the baseline firing rates of each neuron population in the absence of external inputs, according to Eshel et al. (2015) experimental data - with external inputs, the baseline activity of DA neurons is around 5 Hz.

The parameter *r* sets the balance of α4β2 nAChR action through GABA or DA neurons in the VTA. For *r* = 0, they act through GABA neurons only, whereas for *r* = 1 they influence DA neurons only. Φ(.) is the linear rectifier function, which only keeps the positive part of the operand and outputs 0 when it is negative. *F* (.) is a non-linear sigmoid transfer function for the dopaminergic neurons enabling to describe the high firing rates in the bursting mode and the low frequency activity in the tonic (pacemaker) mode, and their slow variation below their baseline activity with external inputs (≃5 Hz):

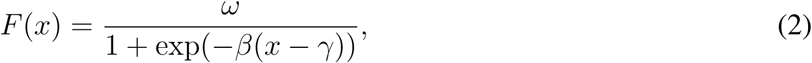

where *ω* = 30 represent the maximum firing rate, *γ* = 8 is the inflexion point and *β* = 0.3 is the slope. These parameters were chosen in order to account for bursting activity of DA neurons starting from a certain threshold (*γ*) of input and their maximal activity observed *in vivo* (Hyland et al., 2002; Eshel et al., 2015). Indeed, physiologically, high firing rates (> 8 Hz) are only attained during DA bursting activity and not tonic activity (≃ 5 Hz).

The input currents in Eq. 1 are given by:

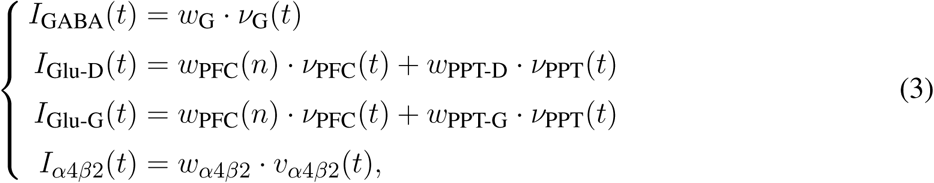

where *w*_x_’s (with x = G, PFC, PPT-D, PPT-G, α4β2) specify the total strength of the respective input (Fig. 1, Table 1). The weight of α4β2-nAChRs, *w*_*α*4*β*2_ = 15 was chosen in order to account for the increase of baseline firing rates compared to (Graupner et al., 2013) where *w*_*α*4*β*2_ = 1, *B*_D_ = 0.1 and *B*_G_ = 0.

Inhibitory input to DA cells, *I*_GABA_, depends on GABA neuron population activity, *ν*_G_ (Eshel et al., 2015). Excitatory input to DA and GABA cells depends on PFC-NAc (Ishikawa et al., 2008; Keiflin and Janak, 2015) and PPTg (Lokwan et al., 1999; Yoo et al., 2017) glutamatergic inputs activities, *ν*_PFC_ and *ν*_PPT_ respectively (see next section). The activation of α4β2 nAChRs, *v*_*α*4*β*2_, determines the level of direct excitatory input *I*_*α*4*β*2_ evoked by nicotine or acetylcholine (see last section).

### 2.2 Neuronal activities during classical-conditioning

As described above, previous studies identified signals from distinct areas that could be responsible for VTA DA neurons activity during classical conditioning. We thus consider a simple model that particularly accounts for Eshel et al. (2015) experimental data on VTA GABA neurons activity. In this approach, we propose that the sustained activity reflecting reward expectation in GABA neurons comes from the PFC (Schoenbaum et al., 1998; Le Merre et al., 2018), that sends projections on both VTA DA and GABA neurons through the NAc (Morita et al., 2013; Keiflin and Janak, 2015). The PFC-NAc pathway thus drives feed-forward inhibition onto DA neurons by exciting VTA GABA neurons that in turn inhibit DA neurons (Fig. 1). Second, we consider that a subpopulation of the PPTg provides the reward signal to the dopamine neurons at the US (Kobayashi and Okada, 2007; Okada et al., 2009).

#### 2.2.1 Classical-conditioning task and the associated signals

We modeled a VTA neural circuit (Fig. 1) while mice are classically conditioned with a tone stimulus that predicts an appetitive outcome as in (Eshel et al., 2015), but with 100% probability. Each simulated behavioral trial begins with a conditioned stimulus (CS; a tone, 0.5 s), followed by an unconditioned stimulus (US; the outcome, 0.5 s) separated by an interval of 1.5 s. (Fig. 2A). This type of task, implying a delay between the CS offset and the US onset (here, 1 s), is then a trace-conditioning task, that differs from a delay-conditioning task where the CS and US overlap (Connor and Gould, 2016).

**Figure 2.**
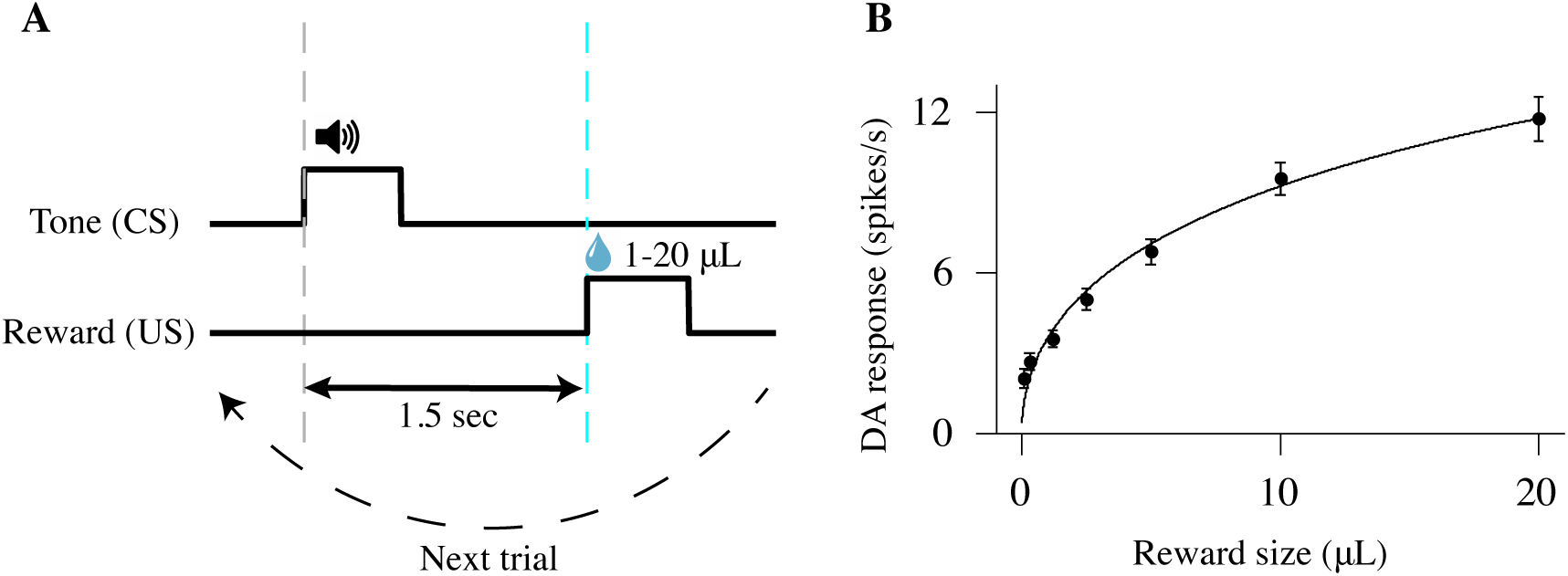
Schematic of a classical-conditioning task (A) Simulated thirsty mice receive a water reward ranging from 1 to 20 μL. Tone (CS) and reward (US) onsets are separated by 1.5 sec. (B) Firing rates (mean ± standard-error (s.e.)) of optogenetically identified dopamine neurons in response to different sizes of unexpected reward. Adapted from (Eshel et al., 2016).

As the animal learns that a fixed reward constantly follows a predictive tone at a specific timing, our model proposes possible underlying biological mechanisms of Pavlovian-conditioning in PPTg, PFC, VTA DA and GABA neurons (Fig. 1).

As represented in previous models (O’Reilly et al., 2007; Vitay and Hamker, 2014), the CS signal is modeled by a square function (*ν*_CS_(*t*)) equal to 1 during the CS presentation (0.5 s) and to 0 otherwise (Fig. 2A). The US signal is modeled by a similar square function (*ν*_US_(*t*)) as the CS but is equal to the reward size during the US presentation (0.5 s) and 0 otherwise (Fig. 2A).

#### 2.2.2 Neural representation of US signal in the PPTg

Dopamine neurons in the VTA exhibit a relatively low tonic activity (around 5 Hz), but respond phasically with a short-latency (< 100 ms), short-duration (< 200 ms) burst of activity in response to unpredicted rewards (Schultz, 1998; Eshel et al., 2015). These phasic bursts of activity are dependent on glutamatergic activation by a subpopulation of PPTg (Okada et al., 2009; Keiflin and Janak, 2015; Yoo et al., 2017) found to discharge phasically at reward delivery, with the levels of activity associated with the actual reward and not affected by reward expectation.

To integrate the US input into a short-term phasic component we use the function *G_τ_* (*x*(*t*)) (Vitay and Hamker, 2014) defined as follows:

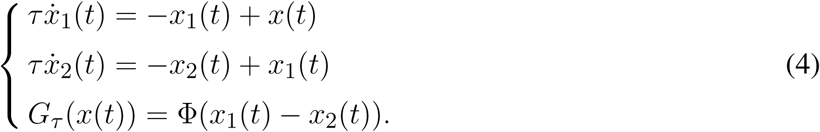

Here when *x*(*t*) switches from 0 to 1 at time *t* = 0, *G_τ_* (*x*(*t*)) will display a localized bump of activation with a maximum at *t* = *τ* . This function is thus convenient to integrate the square signal *ν*_US_(*t*) (Fig. 2A) into a short-latency response.

Furthermore, dopamine response amplitudes to unexpected rewards follow a simple saturating function (fitted by a Hill function in Fig. 2B) (Eshel et al., 2015, 2016). We thus consider that PPTg neurons respond to the reward delivery signal (US) in a same manner as DA neurons *i*.*e*. with a saturating dose-response function:

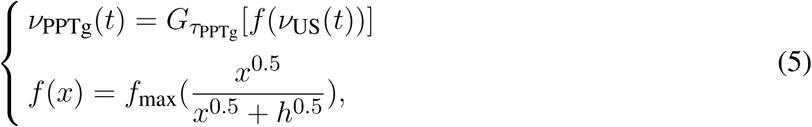

where *ν*_PPTg_ is the mean activity of the PPTg neurons population, *τ*_PPTg_ = 100 ms (the short-latency response), and *f* (*x*) is a Hill function with two parameters: *f*_max_, the saturating firing rate; and *h*, the reward size that elicits half-maximum firing rate. Here, we chose *f*_max_ = 70 and *h* = 20 in order to obtain a similar dose-response curve once PPTg activity is transferred to DA neurons as in (Eshel et al., 2016) (Fig. 2B).

#### 2.2.3 Neural representation of CS signal in the PFC

In addition to their response to unpredicted rewards, DA neurons learn to respond to reward-predictive cues and to reduce their response at the US (Schultz et al., 1997; Schultz, 1998; Matsumoto and Hikosaka, 2009; Eshel et al., 2015). Neurons in the PFC respond to these cues through a sustained activation starting at the CS onset and ending at the reward-delivery (Connor and Gould, 2016; Le Merre et al., 2018). Furthermore, this activity has been shown to increase in the early stage of a classical-conditioning learning task (Schoenbaum et al., 1998; Le Merre et al., 2018). Especially, the PFC participates in the association of temporally separated events in trace-conditioning task through working-memory mechanisms (Connor and Gould, 2016), maintaining a representation of the CS accross the the CS-US interval, and this timing-association is dependent on dopamine modulation in the PFC (Puig et al., 2014; Popescu et al., 2016).

We thus assume that the PFC integrates the CS signal and learns to maintain its activity until the reward delivery. Consistently with previous neural-circuit working-memory models (Durstewitz et al., 2000), we minimally described this mechanism by a neural population with recurrent excitation and a slower adaptation inspired from (Gerstner et al., 2014):

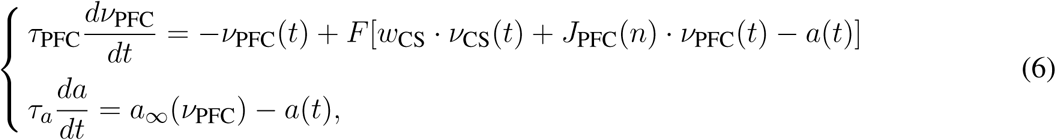

where *τ*_PFC_ = 100 ms (short-latency response), *a*(*t*) describes the amount of adaptation that neurons have accumulated, *a*_∞_ = *c* ⋅ *ν*_PFC_ is the asymptotic level of adaptation that is attained by a slow time constant *τ_a_* = 1000 ms (Gerstner et al., 2014) if the population continuously fires at a contant rate *ν*_PFC_, *J*_PFC_(*n*) represents the strength of the recurrent excitation exerted by the PFC depending on the learning trial *n* (initially *J*(1) = 0.2), *w*_CS_ the strength of the CS input. *F* (*x*) is the non-linear sigmoid transfer function defined in Eq. 2 allowing the emergence of bistability network. We chose *ω* = 30, *γ* = 8 and *β* = 0.5 in order to account for the PFC activity changes in working-memory tasks (Connor and Gould, 2016).

#### 2.2.4 Learning of the US timing in the PFC

The dynamic system described above typically switches between two stables states: quasi absence of activity or maximal activity in the PFC. The latter stable state particularly appears as *J*_PFC_(*n*) increases with learning:

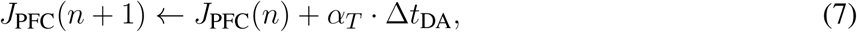

where *α_T_* = 0.2 is the timing learning rate, Δ*t*_DA_ = *t*_2_ – *t*_1_ measures the difference between the time at which PFC activity declines (*t*_1_ such as *ν*_PFC_(*t*_1_) ≃*γ* after CS onset) and the time of DA maximal activity at the US, *t*_2_. This learning mechanism of reward timing, simplified from Luzardo et al. (2013), triggers the increase of the recurrent connections (*J*_PFC_) through dopamine-mediated modulation in the PFC (Puig et al., 2014; Popescu et al., 2016) such as *ν*_PFC_ collapses at the time of reward delivery. This learning process occurs in the early stage of the task (Le Merre et al., 2018) and is therefore much faster than the learning of reward expectation.

#### 2.2.5 Learning of reward expectation in cortico-striatal connections

According to studies showing a DA-dependent cortico-striatal plasticity (Yagishita et al., 2014; Keiflin and Janak, 2015), we assumed that the reward value predicted from the tone (CS) is stored in the strength of cortico-striatal connections (*w*_PFC_(*n*)), *i*.*e*. between the PFC and the NAc, and is updated through plasticity mechanisms depending on phasic dopamine response after reward delivery as in the following equation proposed by (Morita et al., 2013):

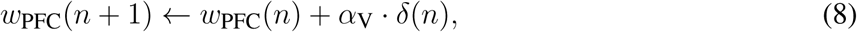

where *α*_V_ is the cortico-striatal plasticity learning rate related to reward magnitude, *δ*(*n*) is a deviation from the DA baseline firing rate, computed by the area under curve of *ν*_D_ in a 200 ms time-window following US onset, above a baseline defined by the value of *ν*_D_ at the time of US onset. *δ*(*n*) is thus the reward-prediction error signal that updates the reward-expectation signal stored in the strength of the PFC input *w*_PFC_(*n*) until the value of the reward is learned (Rescorla and Wagner, 1972).

This assumption was taken from Morita et al. (2013) modeling work and various hypotheses on dopamine-mediated plasticity in associative-learning (Keiflin and Janak, 2015) and recent experimental data (Yagishita et al., 2014; Fisher et al., 2017). It implies that the excitatory signal from the PFC first activates the nucleus accumbens (NAc) and is then transferred via the direct excitatory pathway to the VTA. Here, we then considered that *w*_PFC_ is provided by the PFC-NAc pathway but we did not explicitly represent the NAc population(Fig. 1).

#### 2.2.6 Cholinergic input activity

Our model also reflects the cholinergic (ACh) afferents to the DA and GABA cells in the VTA (Dautan et al., 2016; Yau et al., 2016). The α4β2 nAChRs are placed somatically on both the DA and the GABA neurons and their activity depends on ACh and Nic concentration within the VTA (see last section). As PPTg was found to be the main source of cholinergic input to the VTA, we assume that ACh concentration directly depends on PPTg activity, as modeled by the following equation:

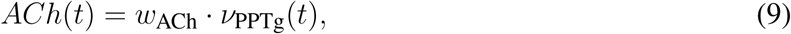

where *w*_ACh_ = 1 μM is the amplitude of the cholinergic connection that tunes concentration of acetylcholine *ACh* (in μM) at a physiologically relevant concentration (Graupner et al., 2013).

### 2.3 Modeling the activation and desensitization of nAChRs

We implemented nAChR activation and desensitization from (Graupner et al., 2013) as transitions of two independent state variables: an activation gate and a desensitization gate. The nAChR receptors can then be in four different states: deactivated/sensitized, activated/sensitized, activated/desensitized and deactivated/desensitized. The receptors are activated in response to both Nic and ACh, while desensitization is driven by Nic only (if *η* = 0). Once Nic or ACh is removed, the receptors can switch from activated to deactivated and from desensitized to sensitized.

The mean total activation level of nAChRs (*ν*_*α*4*β*2_) is modeled as the product of the activation rate *a* (fraction of receptors in the activated state) and the sensitization rate *s* (fraction of receptors in the sensitized state). The total normalized nAChR activation is therefore: *ν*_*α*4*β*2_ = *a* ⋅ *s*. The time course of the activation and the sensitization variables is given by:

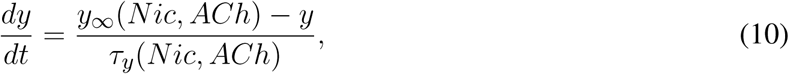

where *τ_y_* (*Nic*, *ACh*) refers to the Nic/ACh concentration-dependent time constant at which the steady-state *y*_∞_(*Nic*, *ACh*) is achieved. The maximal achievable activation or sensitization, for a given Nic/ACh concentration, *a*_∞_(*Nic*, *ACh*) and *s*_∞_(*Nic*, *ACh*) are given by Hill equations of the form:

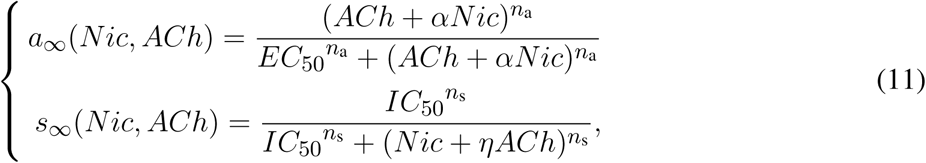

where *EC*_50_ and *IC*_50_ are the half-maximal concentrations of nAChR activation and sensitization, respectively. The factor *α* > 1 accounts for the higher potency of Nic to evoke a response as compared to ACh: *α*_*α*4*β*2_ = 3. *n*_a_ and *n*_s_ are the Hill coefficients of activation and sensitization. *η* varies between 0 and 1 and controls the fraction of the ACh concentration driving receptor desensitization. Here, as we only consider Nic-induced desensitization, we set *η* = 0.

As the transition from the deactivated to the activated state is fast (^~^μs), the activation time constant *τ_a_* was simplified to be independent on ACh and Nic concentration: *τ_a_*(*Nic*, *ACh*) = *τ_a_* = *const*. The time course of Nic-driven desensitization is characterized by a concentration-dependent time constant

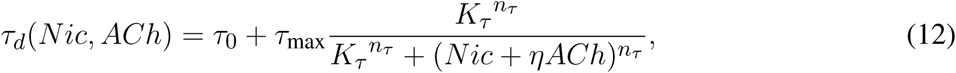

where *τ*_max_ refers to the recovery time constant from desensitization in the absence of ligands, *τ*_0_ is the fastest time constant at which the receptor is driven into the desensitized state at high ligand concentrations. *K_τ_* is the concentration at which the desensitization time constant attains half of its minimum. All model assumptions are further described in Graupner et al. (2013).

### 2.4 Simulated experiments

#### 2.4.1 Optogenetic inhibition of VTA GABA neurons

In order to qualitatively reproduce Eshel et al. (2015) experimental data, we simulated the photo-inhibition effect in a subpopulation of VTA GABA neurons with an exponential decrease between *t* = 1.5 s and *t* = 2.5s (±500 ms around reward-delivery). First, the light was modeled by a square signal *ν*_light_ equal to the laser intensity *I* = 4 for 1.5 < *t* < 2.5 and zero otherwise. Then, we subtracted this signal to VTA GABA neuron activity as follows:

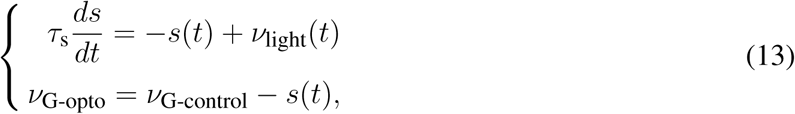

where *s* is the subtracted signal that integrates the light signal *ν*_light_ with a time constant *τ*_s_ = 300 ms, *ν*_G-opto_ is the photo-inhibited GABA neurons activity, and *ν*_G-control_ is the normal GABA neurons activity with no opto-inhibition. All parameters (*I*, *τ*_s_) were chosen in order to reproduce qualitatively the photo-inhibition effects revealed by Eshel et al. (2015) experiments (Fig. **??**C). Furthermore, as the effects of GABA photo-inhibition onto DA neurons appear to be relatively weak (Fig. **??**D, green trace), we assumed that only a subpopulation of the total GABA neurons are photo-inhibited and we therefore applied Eq. 13 for only 20% of the VTA GABA population. This assumption was based on the partial expression of Archeorhodopsin (ArchT) in GABA neurons (Eshel et al. (2015), Extended Data Fig. 1) and the other possible optogenetic effects (recording distance, variability of the response among the population, laser intensity, etc.).

#### 2.4.2 Nicotine injection in the VTA

In order to model chronic nicotine injection in the VTA while mice perform classical-conditioning tasks with water reward, the above equations were simulated but after 5 min of 1 μM Nic injection in the model for each trial. This process allowed to focus only on the effects of α4β2-nAChRs desensitization (see next section) during conditioning trials.

#### 2.4.3 Decision-making task

We simulated a protocol designed by Naudé et al. (2016) recording simultaneously the sequential choices of a mouse between three differently rewarding locations (associated with reward size) in a circular open-field (Fig. 7A). These three locations form an equilateral triangle and provide respectively 2, 4, 8 μL water rewards. Each time the mouse reaches one of the rewarding locations, the reward is delivered. However, the mouse receives the reward only when it alternates between rewarding locations.

Before the simulated task, we considered that the mouse has already learned the value of each location (pre-training) and thus knows the expected associated reward. Each value was computed taking the maximal activity of DA neurons within a time window following the CS onset (here, the view of the location) for the three different reward sizes after learning. We also considered that each time the mouse reaches a new location, it enters in a new state *i*. Decision making-models inspired from Naudé et al. (2016) determine the probability *P_i_* of choosing the next state *i* as a function of the expected value of this state. Because mice could not return to the same rewarding location, they had to choose between the two remaining locations. We thus modeled decisions between two alternatives. The probability *P_i_* was computed according to the softmax choice rule:

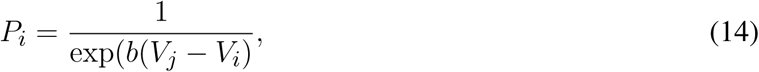

where *V_i_* and *V_j_* are the values of the states *i* and *j* (the other option) respectively, *b* is an inverse temperature parameter reflecting the sensitivity of choice to the difference between both values. We chose *b* = 0.4 which corresponds to a reasonable exploration-exploitation ratio.

We simulated the task over 10,000 simulations and computed the number of times the mouse chose each location. We thus obtained the average repartition of the mouse over the three locations. A similar task was simulated for mice after 5 min Nic ingestion (see below).

## 3 Results

We used the model developed above to understand the learning dynamics within the PFC-VTA circuitry and the mechanisms by which the RPE in the VTA is constructed. Our minimal circuit dynamics model of the VTA was inspired from Graupner et al. (2013) and modified according to recent neurobiological studies (see Methods) in order to reproduce RPE computations in the VTA. This model reflects the glutamatergic (from PFC and PPTg) and cholinergic (from PPTg) afferents to VTA DA and GABA neurons, as well as local inhibition of DA neurons to GABA neurons. We also included the activation and desensitization dynamics of α4β2 nAChRs from (Graupner et al., 2013), placed somatically on both DA and GABA neurons, depending on a fraction parameter *r*. We simulated the proposed PFC and PPTg activity during the task, where corticostriatal connections between the PFC and the VTA and recurrent connections among the PFC were gradually modified by dopamine in the NAc. Finally, we studied the potential influence of nicotine exposure on DA responses to rewarding events.

We should note that most experiments we simulated herein concern the learning task of a CS-US association (Fig. 2). The learning procedure consists of a conditioning phase where a tone (CS) and a constant water-reward (US) are presented together for 50 trials. Within each 3 s-trial, the CS is presented at *t* = 0.5 s (Fig. 3, 5, 6, dashed grey line) followed by the US at *t* = 2 s (Fig. 3, 5, 6, dashed cyan line).

**Figure 3.**
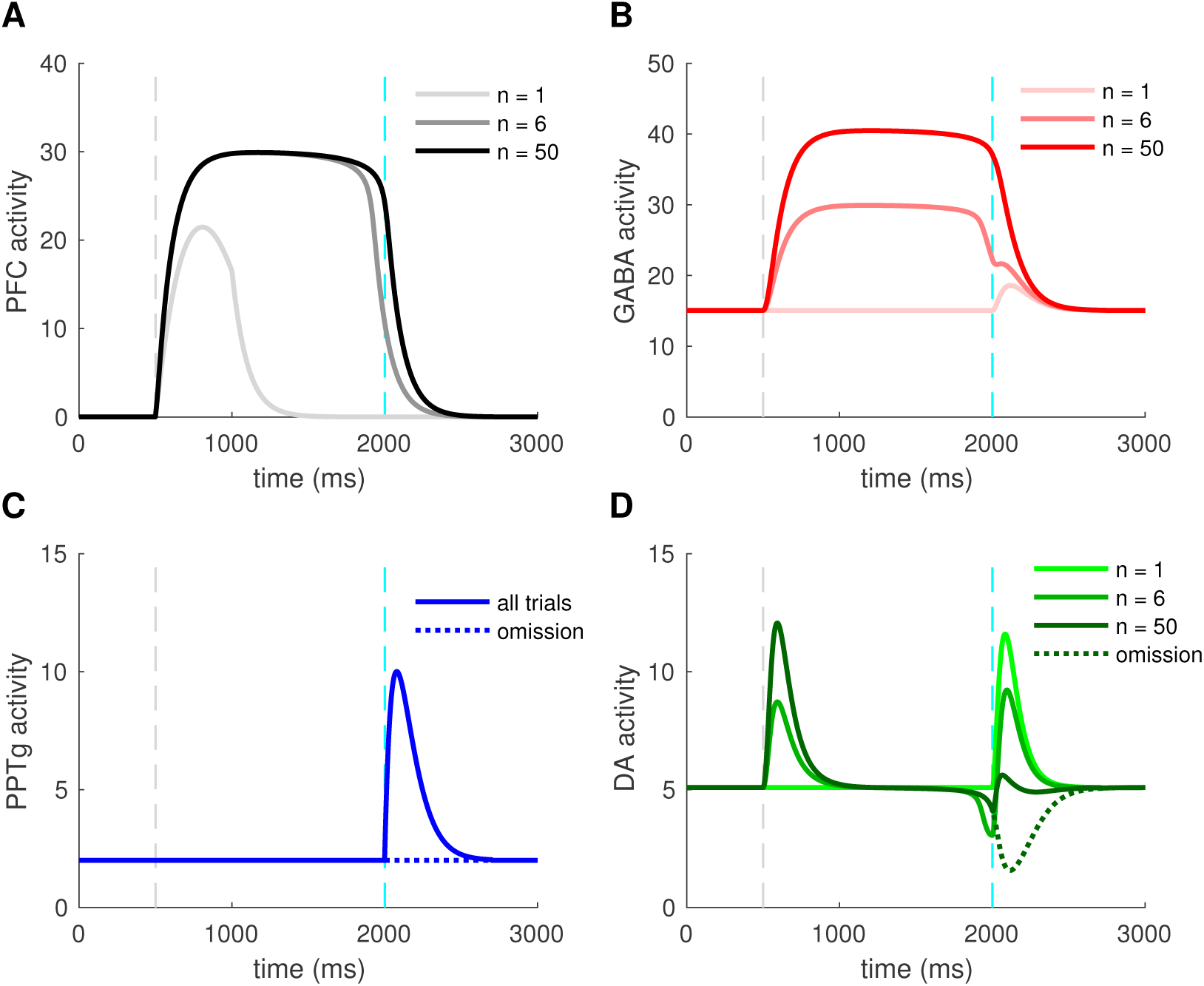
Activity of VTA neurons and their afferents during a pavlovian-conditioning task Simulated mean activity (Hz) of each neuron population during a pavlovian-conditioning task, where a tone is presented systematically 1.5 s before a water reward (4 μL). Three different trials are represented: the initial conditioning trial (*n* = 1, light colors), an intermediate trial (*n* = 6, medium colors) and the final trial (*n* = 50, dark colors) and when reward is omitted after learning (dotted lines). Vertical dashed grey and cyan lines represent CS and US onsets, respectively. (A) PFC neurons learn the timing of the task by maintaining their activity until US. (C) PPTg neurons activity responds to the US signal at all trials.(B) VTA GABA persistent activity increases with learning, (D) VTA DA activity increase at the CS and decrease at the US.

### 3.1 Pavlovian-conditioning task and VTA activity

DA activity during a classical-conditioning task was first recorded by Schultz (1998) and tested in further several studies. Additionally, Eshel et al. (2015) also recorded the activity of their putative neighbor neurons, the VTA GABA neuron population. Our goal was first to qualitatively reproduce VTA GABA and DA activity during associative learning of a pavlovian-conditioning task.

In order to understand how different brain areas interact during conditioning and reward omission, we examined the simulated time course of activity of four populations (PFC, PPTg, VTA DA and GABA), Fig. 3, at the initial conditioning trial (*n* = 1, light color curves), an intermediary trial (*n* = 6, medium color curves) and at the final trial (*n* = 50, dark color curves). In line with experiments, the reward delivery (Fig. 3, dashed cyan lines) activates the PPTg nucleus (Fig. 3C) at each conditioning trial. These neurons activate in turn VTA DA and GABA neurons through glutamatergic connections, causing a phasic burst in DA neurons at the US when the reward is unexpected (Fig. 3D, *n* = 1), and a small excitation in GABA neurons (Fig. 3B, *n* = 1). PPTg fibers also stimulate VTA neurons through ACh-mediated α4β2 nAChRs activation, with a larger influence on GABA neurons (*r* = 0.2 in Fig. 1).

Early in the conditioning task, simulated PFC neurons respond to the tone (Fig. 3A, *n* = 1), and this activity builds up until being maintained during the whole CS-US interval (Fig. 3A, *n* = 6, *n* = 50). Thus, PFC neurons show a working-memory like activity now tuned to decay at the reward delivery time. Concurrently, the phasic activity of DA neurons at the US acts as prediction-error signal on corticostriatal synapses, increasing the glutamatergic input from the NAc onto VTA DA and GABA neurons (Fig. 3B, 3D, 4B). Note that the NAc was not modeled explicitly, but we modeled the net effect of the PFC-NAc plasticity with the variable *w*_PFC_ (see next section).

**Figure 4.**
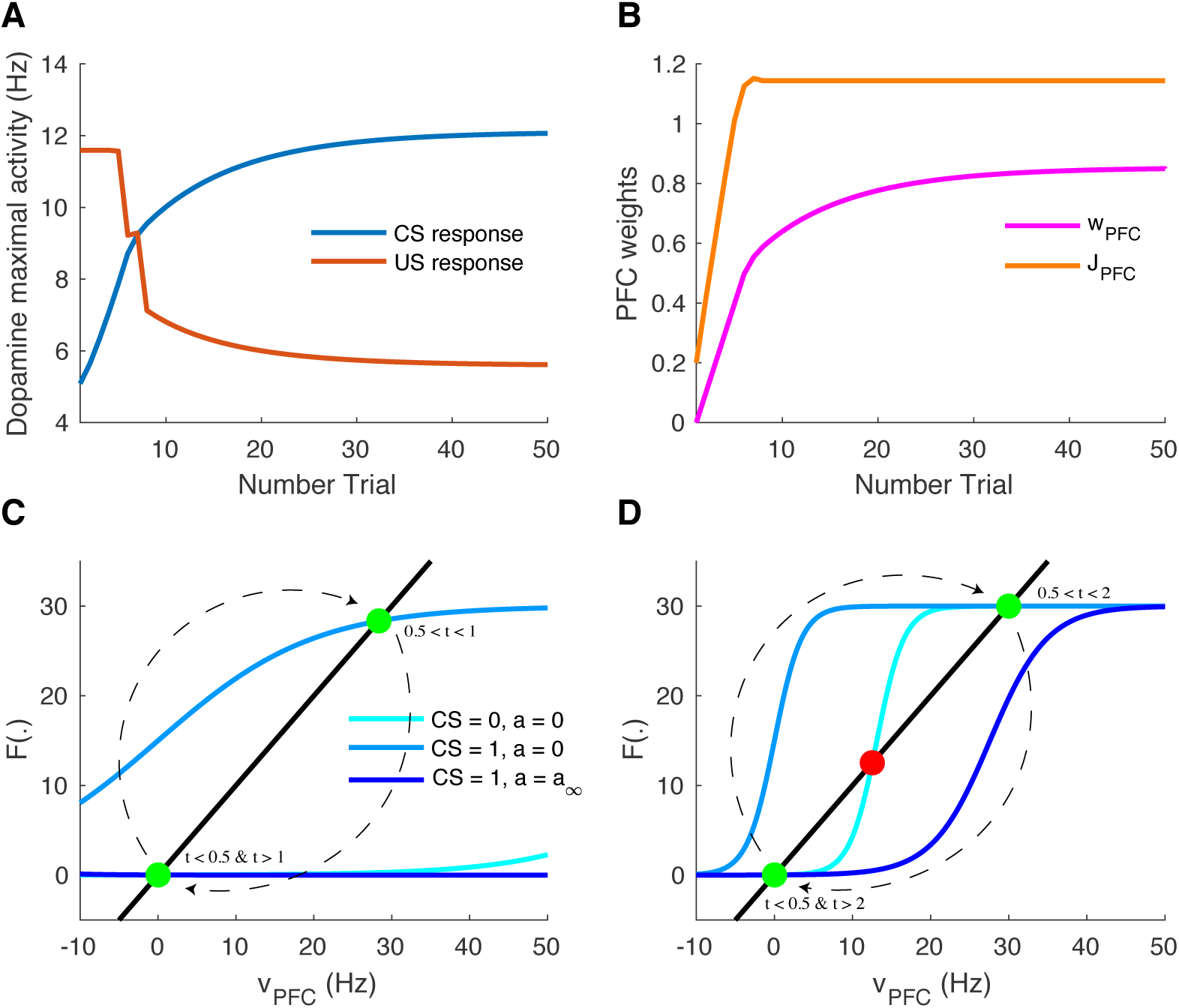
Learning of reward timing and magnitude during classical-conditioning (A) The maximal activity of the VTA DA neurons at the CS onset (blue line) and at the reward delivery (orange line) is plotted for each trial of the conditioning task. These values are computed by taking the maximum value of the firing rate of the DA neurons in a small time window (200 ms) after the CS and the US onsets. (B) PFC weights showing two phases of learning: learning of the US timing by PFC recurrent connections weight (*J*_PFC_, orange line) and learning of the reward value by the weights of PFC neurons onto VTA neurons (*w*_PFC_, magenta line). (C,D) Phase analysis of PFC neuron activity from Eq. 6 before learning (C) and after learning (D). Different times of the task are represented: *t* < 0.5 s (before CS onset, light blue) and 1 s < *t* < 2 s (between CS offset and US onset, light blue), 0.5 s < *t* < 1 s (during CS presentation, medium blue) and *t* > 2 s (after US onset, dark blue). Fixed points are represented by green (stable) or red (unstable) dots. Dashed arrows: trajectories of the system from *t* = 0 to *t* = 3 s.

Consequently, with learning, VTA GABA neurons show a sustained activation during the CS-US interval (Fig. 3B, *n* = 6, *n* = 50) as found in Eshel et al. (2015) experiments and in turn inhibit their neighboring dopamine neurons. Thus, in DA neurons, the GABA neurons-induced inhibition occurs with a slight delay after the PFC-induced excitation, resulting in a phasic excitation at the CS and a phasic inhibition at the US (Fig. 3D, *n* = 50).

The latter inhibition progressively cancels the reward-evoked excitation by the PPTg glutamatergic fibers in DA neurons. It also accounts for the pause in DA firing when reward is omitted after learning (Fig. 3B, 3D, *n* = 50, dashed lines).

Together, these results propose a simple mechanism for RPE computation the VTA and its afferents.

Let us now take a closer look at the evolution of the phasic activity of DA neurons and their PFC-NAc afferents during the conditioning task. Fig. 4A shows the evolution of CS- and US-mediated DA peaks over the 50 conditioning trials. Firstly, the US-related bursts (Fig. 4A, red line) remain constant in the early trials until the timing is learnt by the PFC recurrent connections *J*_PFC_ (Fig. 4B, orange line) following Eq. 7. Secondly, US and CS (Fig. 4A, blue line) responses respectively decrease and increase over all trials, following a slower learning process from cortico-striatal connections (Fig. 4B, magenta line) described by Eq. 8. This two-speed learning process enables to qualitatively reproduce the DA dynamics found experimentally, with almost no effect outside the CS and US time-windows (Fig. 3D).

Particularly, the graphical analysis of the PFC system enables us to understand the timing learning mechanism. From Eq. 6, we can see where the two functions *ν*_PFC_ → *ν*_PFC_ and *ν*_PFC_ → *F* [*w*_CS_ ⋅ *ν*_CS_(*t*) + *J* (*n*) ⋅ *ν*_PFC_(*t*) – *a*(*t*)] intersect each other (fixed points analysis) at four different timings during the simulation: before and after the CS presentation (*ν*_CS_ = 0, *a* = 0), during CS presentation (*ν*_CS_ = 1, *a* = 0) and after the reward is delivered (*ν*_CS_ = 0, *a* = *a*_∞_). Before learning, as *J*_PFC_ is weak (Fig. 4C), the system starts at one fixed point (*ν*_PFC_ = 0), then jumps to another stable point during CS presentation (*ν*_PFC_ ≃ 30) and immediately goes back to the initial point (*ν*_PFC_ = 0) after CS presentation (*t* = 1 s) as shown in Fig. 3A. After learning (Fig. 4D), the system initially shows the same dynamics but when the CS is removed, the system is maintained at the second fixed point (30 Hz) until reward delivery (Fig. 3A, *n* = 50) due to its bistability after CS presentation (cyan curve). Finally, with the adaptation dynamics, the PFC activity decays right after reward delivery (Fig. 4D, dark blue). Indeed, through this timing learning mechanism, the strength of the recurrent connections maintains the Up state activity of the PFC exactly until the US timing (Eq. 7). Together, these simulations show a two-speed learning process that enables VTA dopamine neurons to predict the value and the timing of the water reward from PFC plasticity mechanisms.

### 3.2 Photo-inhibition of VTA GABA neurons modulates prediction errors

We next focus specifically on the local VTA neurons interactions during the conditioning task. Particularly, we model the effects of VTA GABA optogenetic inhibition (Fig. 5) revealed by one of Eshel et al. (2015) experiments. First, in order to reproduce similar VTA activities where reward was delivered with 90% probability, we picked the activity of VTA GABA and DA neurons at an intermediary stage of learning (n = 6), where DA neurons still responded at the US. Second, as in Eshel et al. (2015), we simulated GABA photo-inhibition in a time-window (±500 ms) around the reward delivery time (Fig. 5A, green shaded area). Considering that ArchT virus expression was partial in GABA neurons and that optogenetic effects do not account quantitatively for physiological effects, the photo-inhibition was simulated for only 20% of our GABA population. This simulated inhibition resulted in disinhibition of DA activity during laser stimulation (Fig. 5B). If the inhibition was 100% efficient on GABA neurons, we assume that DA neurons would then burst at high frequencies during the whole period of stimulation.

**Figure 5.**
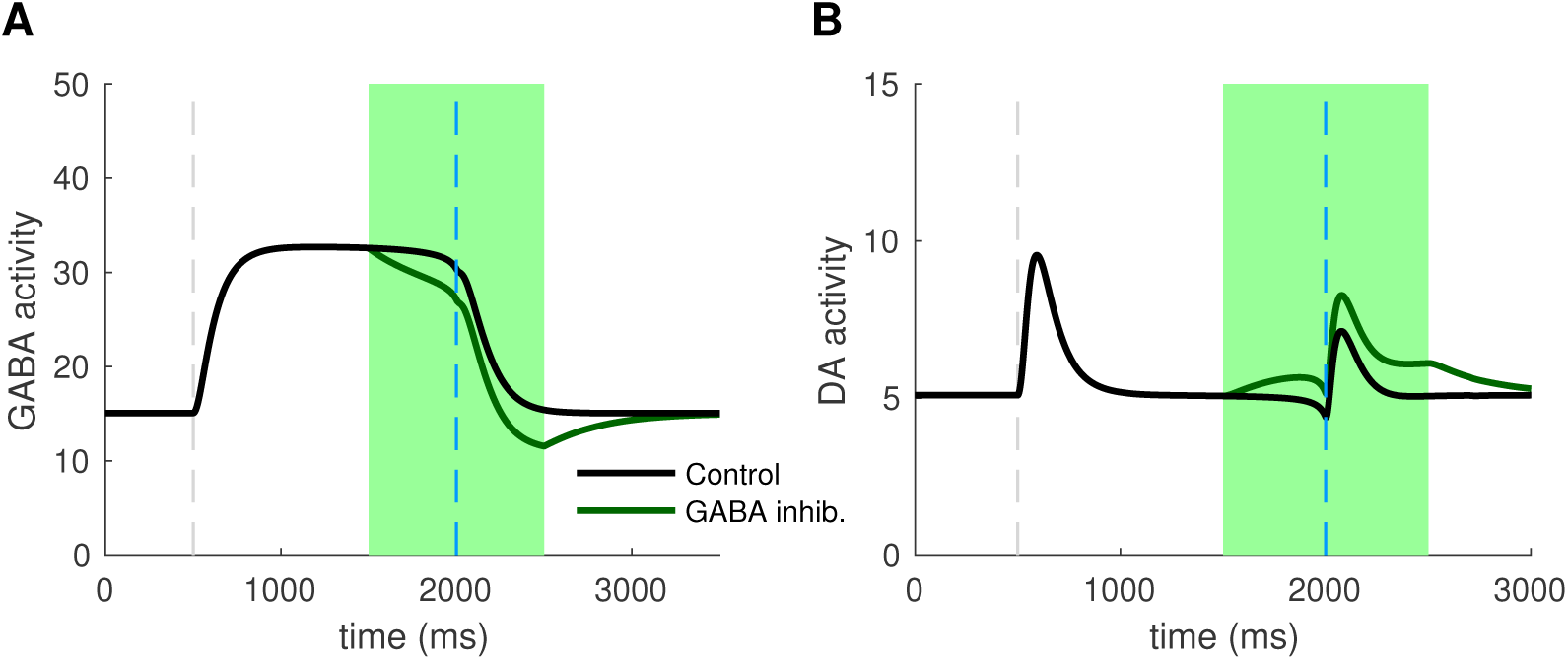
Photo-inhibition of VTA GABA neurons (A) Activity of a subpopulation of GABA neurons (20%) in control (black) and with photo-inhibition (green) simulated by an exponential-like decrease of activity in a ±500 ms time-window around the US (green shaded area). (B) DA activity resulting from GABA neurons activity in control condition (black) and when GABA is photo-inhibited (green).

Inhibiting VTA GABA neurons partially reversed the expectation-dependent reduction of DA response at the US. As proposed by (Eshel et al., 2015), our model accounts for the burst-cancelling expectation signal provided by VTA GABA neurons.

### 3.3 Effects of nicotine on RPE computations in the VTA

We next asked whether we can identify the effects of nicotine action in the VTA during the classical-conditioning task described in Fig. 3. We compared the activity of DA neurons at different conditioning trials to their activity after 5 minutes of 1 μM nicotine injection, corresponding to physiologically relevant concentrations of Nic in the blood after cigarette-smoking (Picciotto et al., 2008; Graupner et al., 2013). For our qualitative investigations, we assume that α4β2-nAChRs are mainly expressed on VTA GABA neurons (*r* = 0.2) and we study the effects of nicotine-induced desensitization on these receptors.

Nic-induced desensitization may potentially lead to several effects. First, under nicotine (Fig. 6B), DA baseline activity slightly increases. Second, simulated exposure also raises DA responses to reward-delivery when the animal is naive (Fig. 6A, 6B, *n* = 1), and therefore to reward-predictive cues when the animal has learnt the task (Fig. 6A, 6B, *n* = 50). As expected, these effects derive from the reduction of the ACh-induced GABA activation provided by the PPTg nucleus (Fig. 3C). Thus, our simulations predict that nicotine would up-regulate DA bursting activity at rewarding events.

**Figure 6.**
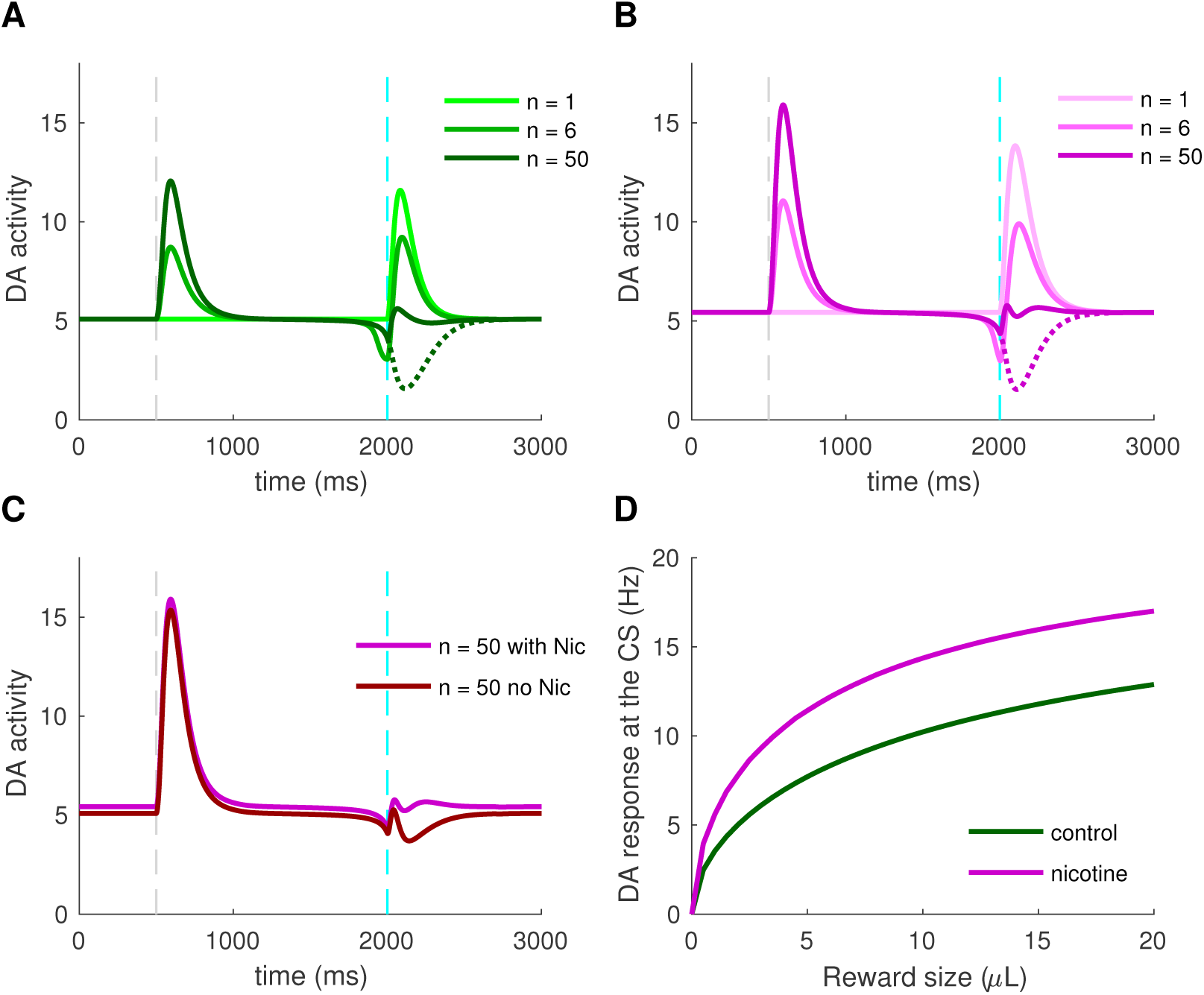
Effects of nicotine on DA activity during classical-conditioning (A) Activity of DA neurons during the pavlovian-conditioning (tone + 4 μL reward) task in three different trials as in Fig. 3. (B) Same as (A) but after 5 min of 1 μM nicotine injection during all conditioning trials. (C) DA activity after learning under nicotine (magenta) or in the same condition but when nicotine is removed (dark red).(D) Dose-response curves of CS-related burst in DA neurons after learning in control condition (green) or under nicotine (magenta).

What would happen if the animal, after having learned in the presence of nicotine, is not exposed to it anymore (nicotine withdrawal)? To answer this question, we investigate the effects of nicotine withdrawal on DA activity after the animal has learnt the CS-US association under nicotine (Fig. 6C), with the same amount of reward (4 μL). In addition to a slight decrease in DA baseline activity, the DA response to the simulated water reward is reduced even below baseline (Fig. 6C, dark red). DA neurons would then signal a negative reward-prediction error, consequently encoding a possible perceived insufficiency of the actual reward it usually receives. From these simulations, we could predict the effect of nicotine injection on the dose-response curve of DA neurons to rewarding events. Here, instead of plotting DA neuron response to different sizes of unexpected rewards as in Fig. 2B, we plot DA response to the CS after the animal has learnt different sizes of rewards, taking the maximum activity in a 200 ms time-window following the CS onset (Fig. 6A, 6B, dark colors). Here, when the animal learns under nicotine, the dose-response curve is elevated, assigning an amplification effect of nicotine on dopamine reward-prediction computations. Notably, the nicotine-induced increase in CS-related bursts grows with the increase of reward size for rewards ranging from 0 to 8 μL. Associating CS amplitude to the predicted value (Rescorla and Wagner, 1972; Schultz, 1998), this suggests that nicotine could increase the value of the cues predicting large rewards, therefore increasing the probability of choosing the associated states compared to control conditions.

### 3.4 Model-based analysis of mouse decision-making under nicotine

In order to evaluate the effects of nicotine on the choice preferences among reward sizes, we simulated a decision-making task where a mouse chose between three locations providing different reward sizes (2, 4, 8 μL) in a circular open-field (Fig. 7A) inspired by Naudé et al. (2016) experimental paradigm.

**Figure 7.**
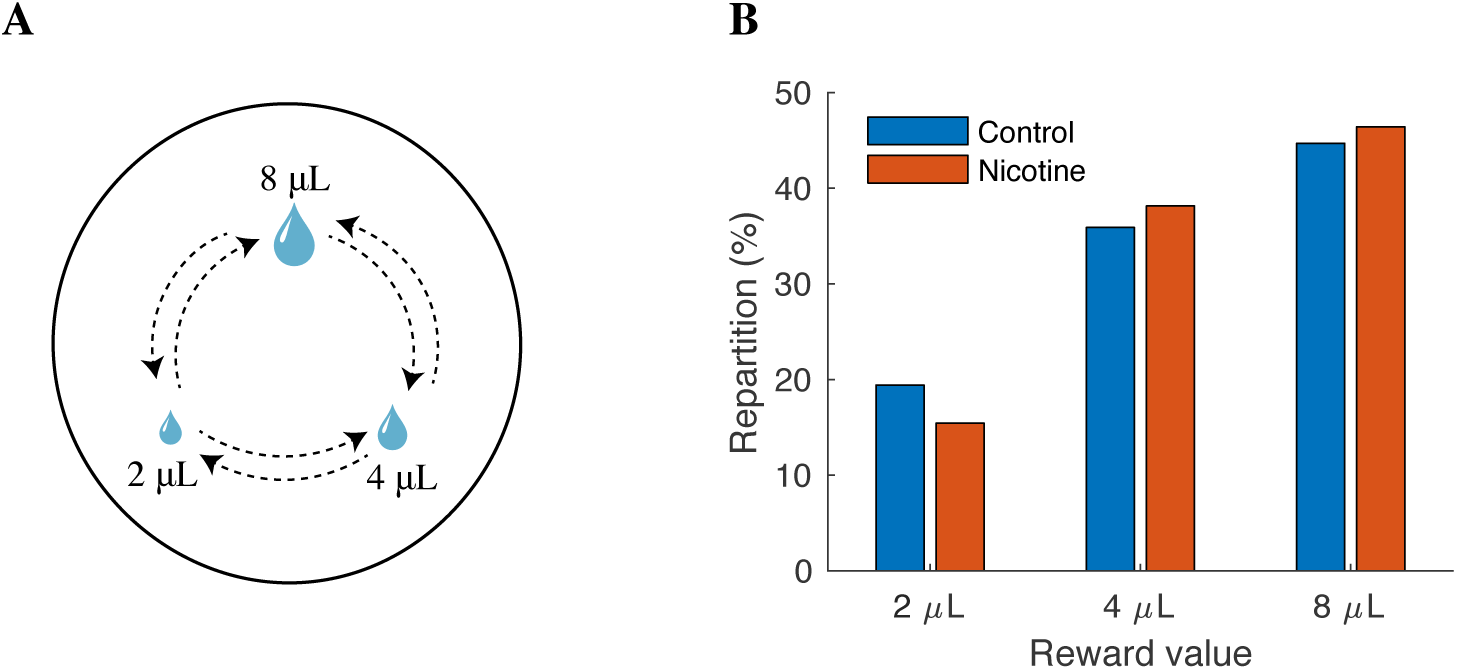
Effects of nicotine on mouse decision-making among reward sizes (A) Illustration of the modeling of the task. Three explicit locations are placed in an open field. Mice receive a reward each time they reach one of the locations. Simulated mice, who could not receive two consecutive rewards at the same location, alternate between rewarding locations. The probability of transition from one state to another depends on the two available options. (B) Proportion of choices of the three rewarding locations as a function of reward value (2, 4, 8 μL) over 10,000 simulations in control mice (blue) or nicotine-ingested mice (red).

Following reinforcement-learning theory (Rescorla and Wagner, 1972; Sutton and Barto, 1998), CS response to each reward size (computed from Fig. 6D) was attributed to the expected value of each location. We then computed the repartition of the mouse between the three locations over 10,000 simulations in control conditions or after 5 min nicotine ingestion.

In control conditions, the simulated mice chose according to the location’s estimated value (Fig. 7B); the mice chose preferentially the locations that provide the greater amount of reward. Interestingly, under Nic-induced nAChRs desensitization, the simulations show a bias of mice choices towards large reward sizes; the proportion of choices for the small reward (2 μL) diminished by about 4%. Thus, these simulations suggested a differential amplifying effect of nicotine for large water rewards.

## 4 Discussion

The overarching aim of this study was to determine how dopamine neurons compute key quantities such as reward-prediction errors, and how these computations are affected by nicotine. In order to do so, we have developed a computational modeling approach extending the population activity of the VTA and its main afferents during a simple task of Pavlovian-conditioning. Including both theoretical and phenomenological conceptions, this model qualitatively reproduces several observations on the VTA activity during the task: phasic DA activity at the US and the CS and persistent activity of VTA GABA neurons. It particularly proposes a two-speed learning process of the reward timing and size mediated by the PFC working memory, coupled with the signaling of reward occurence in the PPTg. Finally, using acetylcholine dynamics coupled with the desensitization kinetics of α4β2-nAChRs in the VTA, we revealed a potential effect of nicotine action on reward perception through up-regulation of DA phasic activity.

### 4.1 Relationship to other computational models

Multiple studies have proposed a dual-pathway mechanism for RPE computation in the brain (O’Reilly et al., 2007; Vitay and Hamker, 2014) through phenomenological bottom-up approaches. Although they propose different possible mechanisms, they mainly gather several components: regions that encode reward-expectation at the CS, regions that encode actual reward, regions that inhibit dopamine activity at the US, and final subtraction of these inputs at the VTA level. These models usually manage to reproduce the key properties of dopamine-related reward activity: progressive appearance of DA bursts at the CS onset, progressive decrease of DA bursts the reward-delivery, phasic inhibition when reward is omitted and early delivery of reward.

Additionally, a top-down theoretical approach as the temporal difference (TD) learning model assumes that the cue and reward cancellation signal both emerge from the same inputs (Sutton and Barto, 1998; Morita et al., 2013). After the task is learned, two sustained expectation signals *V* (*t*) and *V* (*t* + 1) subtract each other (Fig. 8), leading to the TD error: *δ* = *r* + *V* (*t* + 1) – *V* (*t*). Notably, the temporary shift between both signals induce a phasic excitation at CS and an inhibition at the US.

**Figure 8.**
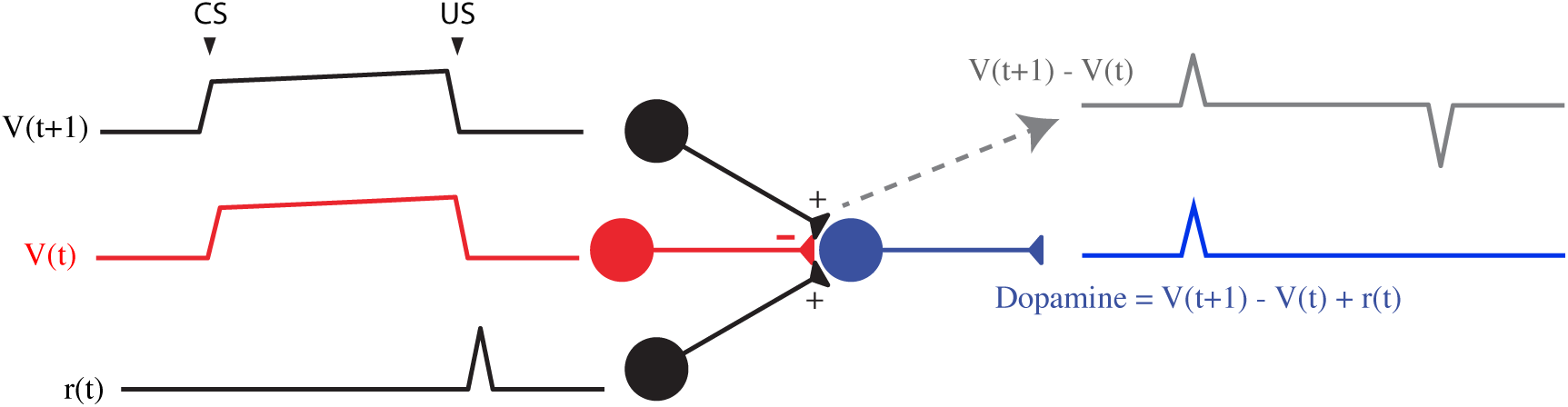
TD learning model (Watabe-Uchida et al., 2017) TD error model as implemented in (Schultz, 1998). The TD error in DA neurons iscomputed from 3 inputs: two reward expectation signals and one reward signal. Traces show how these terms change with time at the last trial of a conditioning task. DA response to a reward omission can be approximated by *V*(*t* + 1) – *V* (*t*) (gray), the derivative of the value function, *V* (*t*). Adapted from (Watabe-Uchida et al., 2017).

TD models are reliable to describe many features of dopamine phasic activity and establish a link between reinforcement learning theory and dopamine activity. However, the biological evidence for such specific signals is still unclear.

In our study, we combine these two phenomenological and theoretical approaches to describe the VTA DA activity. Firstly, our simple model relies on neurobiological mechanisms such as PFC working memory activity (Connor and Gould, 2016; Le Merre et al., 2018), PPTg activity (Kobayashi and Okada, 2007; Okada et al., 2009) and mostly VTA GABA neurons activity (Cohen et al., 2012; Eshel et al., 2015) and describe how these inputs could converge to VTA DA neurons. Secondly, at least at the end of learning, we also proposed a similar integration of inputs as in TD models, with two sustained signals that are temporally delayed. Indeed, the reward expectation signal comes from the same input (PFC): based on recent data on local circuitry in the VTA (Eshel et al., 2015), we assumed that the PFC sends the *V* (*t* + 1) sustained signal to both VTA GABA and DA neurons. Only, via a feed-forward inhibition mechanism, this signal is shifted by VTA GABA neurons membrane time constant *τ*_G_. Thus, in addition to the direct *V* (*t* + 1) excitatory signal from the PFC, VTA GABA neurons would send the *V* (*t*) inhibitory signal to VTA DA neurons (Fig. 8). Adding the reward signal *r*(*t*) provided by the PPTg, our model integrates the TD error *δ* into DA neurons. However, in our model, and as shown in several studies, CS- and US-related bursts gradually increase and decrease with learning, respectively, whereas TD learning predicts a progressive backward shift of the US-related burst during learning, what is not experimentally observed.

Although we make strong assumptions on VTA reward information integration that may be questioned at the level of detailed biology, it proposes a way to explain how the sustained activity in GABA neurons cancel the US-related dopamine burst without affecting the preceding tonic activity of DA neurons during the CS-US interval. Furthermore, this assumption can be strengthened by our simulation of optogenetic experiment (Fig. 5) qualitatively reproducing DA increase in both baseline and phasic activity as found in (Eshel et al., 2015).

### 4.2 Reliability of the VTA afferents

As described above, our model includes two glutamatergic and one GABAergic input to the dopamine neurons, without considering the influence of all other brain areas.

Although the PFC, the NAc and the PPTg were found to be important excitatory afferents to the VTA, it remains elusive whether these signals: 1) respectively encode reward expectation and actual reward and 2) are the only excitatory inputs to the VTA during a classical-conditioning task. As well, it is still unclear whether VTA GABA fully inhibit their dopamine neighbors. Here, we assumed that the activity of DA neurons with no GABAergic input was relatively high (*B*_D_ = 18 Hz) in order to compensate the observed high baseline activity of GABA neurons (*B*_G_ = 14 Hz) and get the observed DA tonic firing rate (≃ 5 Hz). This brings up two issues: do these GABA neurons only partially inhibit their dopamine neighbors, for example, just when activated above their baseline? And also, is the inhibitory reward expectation signal mediated by other brain structures as the LHb (Watabe-Uchida et al., 2012; Tian and Uchida, 2015; Keiflin and Janak, 2015)?

In an attempt to answer this question, Tian et al. (2016) recorded extracellular activity of monosynaptic inputs to dopamine neurons in seven input areas including the PPTg. Showing that many VTA inputs were affected by both CS and US signals, they proposed that DA neurons receive a mix of redundant information and compute a pure RPE signal. However, this does not elucidate which of these inputs effectively affect DA neurons activity during a classical-conditioning task.

While other areas might be implied in RPE computations in the VTA, within our minimal model, we used functional relevant inputs to the VTA that were shown to be strongly affected by reward information based on diverse recurrent studies in the last decades: the working-memory activity in the PFC integrating the timing of reward occurence (Durstewitz et al., 2000; Connor and Gould, 2016), the dopamine-mediated plasticity in the NAc via dopamine receptors (Morita et al., 2013; Yagishita et al., 2014; Keiflin and Janak, 2015), the PPTg activation at the reward delivery (Okada et al., 2009; Keiflin and Janak, 2015). Notably, in most of our assumptions, we rely on experimental data that studied neuronal activity of mice performing a simple classical-conditioning task (reward delivery following conditioning cue with no instrumental actions required). In line with this modeling approach, further optogenetic manipulations implying photo-inhibition as in Eshel et al. (2015) would then be required to study the exact functional impact of the PFC, the NAc and the PPTg on dopamine RPE computations during a simple classical conditioning task.

### 4.3 Nicotine-induced nAChRs desensitization and environmental rewards

Desensitization of α4β2-nAChRs on VTA GABA neurons following nicotine exposure results in increased activity of VTA DA neurons (Mansvelder et al., 2002; Picciotto et al., 2008; Graupner et al., 2013). Through the associative-learning mechanism suggested by our model, nicotine exposure would therefore up-regulate DA-response to rewarding events by decreasing the impact of endogenous acetylcholine on VTA GABA neurons provided by the PPTg nucleus activation (Fig. 6). Together, our results propose that nicotine-mediated nAChRs desensitization potentially enhances the DA response to environmental cues encountered by a smoker (Picciotto et al., 2008).

Indeed, here, we considered that the rewarding effects of nicotine could be purely contextual: nicotine ingestion does not induce a short rewarding stimulus (US), but an internal state (here, after 5 min of ingestion) that would up-regulate smoker perception of environmental rewards (the taste of coffee) and consequently, when learned, the associated predictive cues (the view of a cup of coffee). While nicotine self-administration experiments considered nAChRs activation as the main rewarding effect of nicotine (Picciotto et al., 2008; Changeux, 2010; Faure et al., 2014), our model focuses on the long-term (min to hours) effects of nicotine that a smoker usually seeks, that interestingly correlates with desensitization kinetics of α4β2-nAChRs (Changeux, 2010).

However, the disinhibition hypothesis on nicotine effects in the VTA remains debated. Although demonstrated *in vitro* (Mansvelder et al., 2002) and *in silico* (Graupner et al., 2013), it is still not clear whether nicotine-induced nAChRs desensitization preferentially acts on GABA neurons within the VTA *in vivo*. This would depend on the ratio of α4β2-nAChRs expression levels *r* but also on the preferential VTA targets of cholinergic axons from the PPTg. While we gathered both components into the parameter *r*, recent studies found that PPTg-to-VTA cholinergic inputs preferentially target either DA neurons (Dautan et al.,2016)or GABA neurons (Yau et al., 2016). Notably, accounting for the relevance of Yau et al. (2016) experimental conditions - photo-inhibition of PPTg-to-VTA cholinergic input during a Pavlovian-conditioning task - we chose to preferentially express α4β2-nAChRs on GABA neurons (*r* = 0.2).

Finally, in our behavioral simulations of a decision-making task (Fig. 7), we report that nicotine exposure could potentially bias mice choices towards big rewards. Recent recordings from Faure and colleagues (unpublished data) showed a similar effect of chronic nicotine exposure, with mice showing increasing choices for locations with 100% and 50% reward probabilities at the expense of the location with 25% probability. In this line, future studies could investigate the effects of chronic nicotine on VTA activity during a classical conditioning task as presented here (Fig. 6) but also on behavioral choices according to reward size (Fig. 7).

The idea that dopamine neurons signal reward-prediction errors has revolutionized the neuronal interpretation of cognitive functions such as reward processing and decision-making. While our qualitative investigations are based on a minimal neuronal circuit dynamics model, our results suggest areas for future theoretical and experimental work that could potentially forge stronger links between dopamine, nicotine, learning, and drug-addiction.

The parameters in the model were chosen qualitatively in order to account for most of experimental data from different studies (references) with relative accuracy. The α4β2-containing nAChR parameters were directly taken from (Graupner et al., 2013), whereas the network parameters were qualitatively adapted from different studies. When no data could be related, some parameters were arbitrarily fixed (here).

## Footnotes

**Author Contributions** N.D.: Designed research, performed research, wrote the manuscript. B.S.G.: Designed research, advised N.D., obtained funding, wrote the manuscript.

